# The phase separated CO_2_-fixing pyrenoid proteome determined by TurboID

**DOI:** 10.1101/2022.12.08.519652

**Authors:** Chun Sing Lau, Adam Dowle, Gavin H. Thomas, Philipp Girr, Luke CM Mackinder

## Abstract

Phase separation underpins many biologically important processes such as RNA metabolism, signaling and CO_2_ fixation. However, determining the composition of a phase separated organelle is often challenging due to their sensitivity to environmental conditions which limits the application of traditional proteomics techniques like organellar purification or affinity purification mass spectrometry to understand their composition. In *Chlamydomonas reinhardtii*, Rubisco is condensed into a crucial phase separated organelle called the pyrenoid that improves photosynthetic performance by supplying Rubisco with elevated concentrations of CO_2_. Here, we developed a TurboID based proximity labeling technique in *Chlamydomonas* chloroplasts, where proximal proteins are labeled by biotin radicals generated from the TurboID-tagged protein. Through the expression of two core pyrenoid components fused with the TurboID tag, we have generated a high confidence pyrenoid proxiome that contains the majority of known pyrenoid proteins plus a number of novel pyrenoid candidates. Fluorescence protein tagging of 8 previously uncharacterized TurboID-identified proteins showed 7 were localized to a range of sub-pyrenoid regions. The resulting proxiome also suggests new secondary functions for the pyrenoid in RNA-associated processes and redox sensitive iron-sulfur cluster metabolism. This developed pipeline opens the possibility of investigating a broad range of biological processes in *Chlamydomonas* especially at a temporally resolved sub-organellar resolution.

## Introduction

Nearly all algae contain a microcompartment in their chloroplast called the pyrenoid, which is estimated to be responsible for ∼30% of global CO2 fixation (Mackinder et al., 2016). The pyrenoid of the model green alga *Chlamydomonas reinhardtii* (*Chlamydomonas* hereafter) is a 1-2 micron biomolecular condensate of the principal CO_2_-fixing enzyme Rubisco. It is formed through liquid-liquid phase separation (LLPS) of Rubisco mediated by Essential Pyrenoid Component 1 (EPYC1), that has 5 evenly spaced Rubisco binding motifs (RBM) interspaced by disordered sequence (He et al., 2020; Mackinder et al., 2016; Freeman Rosenzweig et al., 2017; Wunder et al., 2018). The deletion of EPYC1, or the reciprocal binding site of EPYC1 on Rubisco, abolishes pyrenoid formation. With correct pyrenoid assembly being essential for a functional CO_2_ concentrating mechanism (CCM) (Mackinder et al., 2016) that works to saturate Rubisco with CO2 to minimize energetically costly photorespiration, thereby improving photosynthetic efficiency (Wang et al., 2015; Fei et al., 2022). In the face of growing food security issues, the engineering of a pyrenoid-based CCM into major C3 crop plants such as rice, soybean and wheat is regarded as a promising strategy for yield improvement with its prospect of increasing food production by up to 60% (Ray et al., 2013; Long et al., 2019). Recent work has reconstituted a proto-pyrenoid in the higher plant *Arabidopsis thaliana* (Atkinson et al., 2020). However, additional structural components, such as traversing thylakoid membranes and a CO_2_ diffusion barrier will be required for efficient function (Fei et al., 2022). Many of the proteins underpinning these additional structural requirements are unknown making a deep understanding of the structural organization and molecular function of the pyrenoid critical.

Previous pyrenoid proteomes have been achieved via organelle purification (Zhan et al., 2018; Mackinder et al., 2016) and affinity purification mass spectrometry (APMS) (Mackinder et al., 2017), however both these methods have limitations. Whilst multiple robust methods, like APMS, exist to identify strong protein-protein interactions, the ability to identify weak and transient interactions *in vivo* is limited. At a larger spatial scale, subcellular fractionation followed by protein purification and MS is prone to cross-contamination (Christopher et al., 2021). Biomolecular condensates, like the pyrenoid, fall into a class of subcellular structures that are challenging to accurately determine their proteomes as they are typically dynamic, underpinned by weak and transient interactions that are highly sensitive to small changes in the surrounding environment, can vary considerably in size and are not always clearly spatially defined due to the absence of an encapsulating membrane (Hyman et al., 2014; Choi et al., 2020). The recently developed proximity labeling methods such as APEX2 and TurboID (Lam et al., 2015; Branon et al., 2018) are particularly poised to determine the underpinning transient interactions and proteomes of biomolecular condensates (Bracha et al., 2019). APEX2 and TurboID utilize an enzyme tag that drives biotinylation of neighboring proteins *in vivo*. In APEX2 an engineered ascorbate peroxidase converts biotinphenol to biotin-phenoxyl radicals, while in TurboID an engineered biotin ligase generates biotin-5’ -AMP radicals from Biotin and ATP (Branon et al., 2018; Roux et al., 2012). These labile radicals spontaneously biotinylate the surface exposed residues of proteins in close proximity. This reaction gives rise to a localized biotinylation event that is spatially restricted to 10 - 40 nm (Kim et al., 2014, 2016) by the radical’ s diffusion from the enzyme tag. The *in vivo* biotinylation negates the need to purify proteins in their native association, with the high affinity of the biotin tag to streptavidin beads enabling the removal of background contaminants via harsh wash conditions. This results in the identification of strong, weak and transient interactions, in addition to non-interacting proximal proteins. However, since its development proximity labeling has seen limited application in phase-separated systems (Youn et al., 2018; Zhou and Zou, 2021) and has yet to be established in plastids or the model alga *Chlamydomonas*.

In this study we attempt to identify those proteins that were missed in APMS and pyrenoid purification by developing a pyrenoid-based proximity labeling methodology. Using TurboID-based proximity labeling we identify a complementary and robust pyrenoid “proxiome”. Our pyrenoid proxiome contains the majority of previously known pyrenoid proteins and has identified multiple novel pyrenoid components that show distinct sub-pyrenoid localizations via fluorescence tagging. The ability to identify core proteins involved in pyrenoid phase separation highlights the strength of proximity labeling for investigating biomolecular condensate composition and formation. Furthermore, our method establishes proximity labeling in plastids and the leading model algal system, *Chlamydomonas*.

## Results

### *Development of proximity labeling in* Chlamydomonas reinhardtii

We set out to establish proximity labeling in the LLPS pyrenoid within the chloroplast of *Chlamydomonas* (Figure 1A). TurboID has been established in *Arabidopsis* (Mair and Bergmann, 2022; Zhang et al., 2019) and APEX2 in cyanobacteria (Dahlgren et al., 2021) and diatoms (Turnšek et al., 2021). To determine which approach is best suited for *Chlamydomonas*, we designed constructs to test both APEX2 and TurboID (Supplemental Data Set 1). Expression constructs were designed to be compatible with the *Chlamydomonas* modular cloning (MoClo) framework (Crozet et al., 2018) to enable community adoption and compatibility with a broad range of promoters, terminators and selection markers.

**Figure 1.**
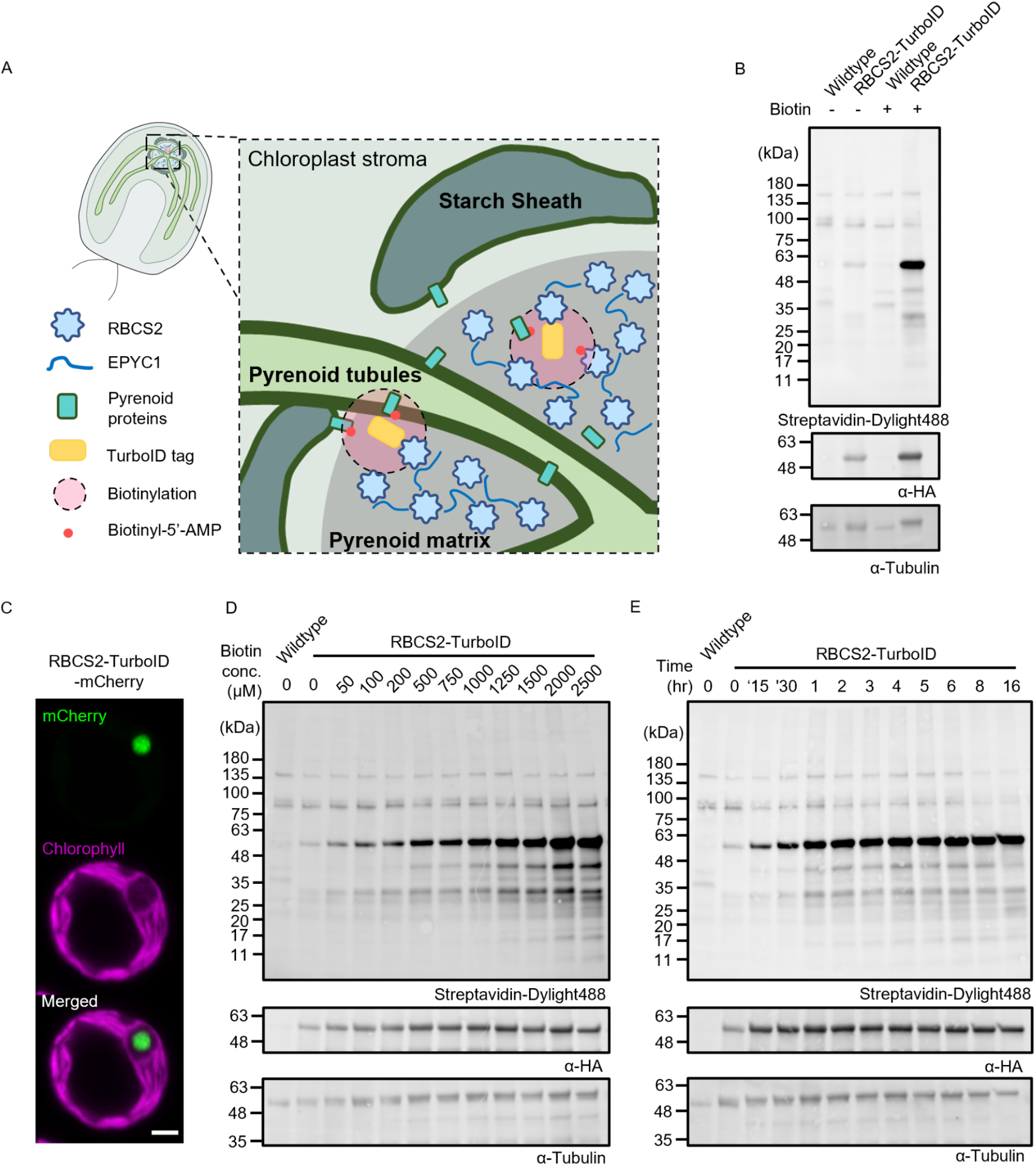
Establishment and optimization of TurboID labeling in the *Chlamydomonas* chloroplast using RBCS2-TurboID lines. **A**, Schematic representation of the *Chlamydomonas* pyrenoid and RBCS2-TurboID. The pyrenoid matrix is surrounded by a starch sheath and traversed by pyrenoid tubules. The initial TurboID construct is targeted to the pyrenoid matrix and expressed as a RBCS2-TurboID fusion protein. Upon addition of biotin substrate short-lived biotin-radicals (red dots) diffuse from the TurboID tag and spontaneously biotinylate neighboring pyrenoid proteins. **B**, Biotinylation signals of lines transformed with the RBCS2-TurboID construct and the untagged background are assessed by immunoblotting whole-cell lysate with a Streptavidin conjugate. Anti-tubulin is used as a loading control, while anti-HA is used to probe for fusion protein expression. **C**, Confocal imaging of RBCS2-TurboID-mCherry. Green and magenta signals represent the mCherry and chlorophyll autofluorescence respectively. Scale bar 2 µm. **D-E**, RBCS2-TurboID labeling efficiency was determined by labeling cells across a biotin concentration gradient (0 - 2500 µM) for 4 hours (**D**) or across a time range (0 - 16 hours) with 2.5 mM Biotin substrate (**E**).

We initially chose the Rubisco small subunit 2 (Cre02.g120150; RBCS2) as our bait due to Rubisco’ s central role in pyrenoid LLPS (Wunder et al., 2018; Meyer et al., 2012), previous data showing that exogenous Rubisco small subunit (RBCS) tagging does not affect CCM functionality (Freeman Rosenzweig et al., 2017), and the availability of known interacting partners for downstream validation (Mackinder et al., 2017; Meyer et al., 2020). Both APEX2 and TurboID were fused to the C-terminus of RBCS2 under the control of the well-established PSAD promoter/terminator pair previously used for fluorescence protein tagging of a broad range of pyrenoid components including RBCS2 (Mackinder et al., 2017). Constructs were transformed via electroporation into the widely used CC-4533 wildtype (WT) strain (Li et al., 2016, 2019). Hygromycin resistant colonies were screened for genomic insertion of the RBCS2 fusion construct via PCR and then for protein expression via immunoblotting against the C-terminal epitope tag (Supplemental Figures 1A and 2A). Expression lines of each construct were named RBCS2-TurboID and RBCS2-APEX2.

We confirmed the correct localization of RBCS2-APEX2 to the pyrenoid by immuno-fluorescence against the 3xFlag tag at the C-terminal of APEX2 (Supplemental Figure 1B). To validate the activity of RBCS2-APEX2, we incubated RBCS2-APEX2 expressing line A2 with biotin-phenol substrate which showed a subtle yet different biotinylation pattern from that of the untagged WT background, especially when activated with higher H2O2 concentration (Supplemental Figure 1C). This led us to pursue a preliminary labeling experiment followed by MS of affinity-purified biotinylated proteins. Analysis of this data showed minimal enrichment for Rubisco or known pyrenoid components (Supplemental Figure 1D). However, when assessing APEX2 peroxidase activity using Amplex red, we detected higher peroxidase activity in RBCS2-APEX2 than the untagged counterpart, suggesting the expressed fusion protein is functional (Supplemental Figure 1E). We tentatively conclude that biotin-phenol has limited cellular permeability resulting in poor labeling. This poor permeability agrees with previous reports in *Saccharomyces cerevisiae*, where cell wall modification was required to facilitate the biotin-phenol uptake (Hwang and Espenshade, 2016; Li et al., 2020). The failure of APEX2 to work in *Chlamydomonas* was also seen in the parallel submission establishing proximity labeling in *Chlamydomonas* (Kreis et al.).

In contrast, initial tests of RBCS2-TurboID showed clear increased biotinylation in comparison to WT with the addition of the biotin substrate (Figure 1B). We observed a pronounced band at ∼50 kDa that likely corresponds to either the self-biotinylation of the RBCS2-TurboID fusion protein (55 kDa) or the Rubisco large subunit (55 kDa) (Figure 1B). A weak biotinylation signal can also be observed in the absence of external biotin addition, indicating that naturally occurring biotin is present in the chloroplast as suggested by the presence of endogenously biotinylated chloroplast proteins (Li-Beisson et al., 2015).

After demonstrating TurboID activity we next assessed the localization of the fusion protein by generating a RBCS2-TurboID-mCherry fusion. Confocal imaging confirmed pyrenoid localization, with the mCherry signal forming a single puncta at the canonical pyrenoid position where there is an absence of chlorophyll signal (Figure 1C). We next optimized biotin substrate concentration and labeling time (Figure 1D and E). Cells were grown photoautotrophically with air-level CO2 supplementation to induce CCM formation that leads to nearly all Rubisco being condensed into the pyrenoid (Borkhsenious et al., 1998). Cells were then incubated with a range of biotin concentrations (0.1 - 2.5 mM) over different time periods (1 - 16 hours). We found that biotin labeling occurs in a substrate (Figure 1D) and time (Figure 1E) dependent manner. In contrast to higher plants where labeling saturation can be achieved by 50 µM biotin (Mair et al., 2019; Wurzinger et al.), labeling in *Chlamydomonas* appears to saturate at a much higher biotin concentration of 2.5 mM. This is in line with work published in parallel in this issue where a 1 mM concentration was used (Kreis et al.). To maximize labeling we performed all later experiments using a final concentration of 2.5 mM biotin. In agreement with previous reports (Zhang et al., 2019; Mair et al., 2019), we similarly observed the rapid activity by TurboID which allowed labeling saturation after ∼1 hour (Figure 1E).

### RBCS2-TurboID labels Rubisco interactors and pyrenoid proteins

We next established a pipeline for streptavidin affinity purification and protein identification by LC-MS/MS (Figure 2A; Methods). Due to the relatively high levels of background biotinylation we set out to further optimize labeling time in a pilot experiment. RBCS2-TurboID and the untagged WT lines were incubated with 2.5 mM biotin across a range of incubation times (1 hr, 2 hr, 4 hr, 8 hr). Proteins extracted from the labeled cells were then subjected to affinity-purification with Streptavidin magnetic beads. A total of 918 proteins were detected by MS across all samples. Initial results showed a strong enrichment for core pyrenoid localized proteins, including RBCS1, RbcL, EPYC1, SAGA2, RBMP1 and RBMP2 when compared to WT cells not expressing RBCS2-TurboID (Figure 2B and Supplemental Data Set 2). Using the detected proteins we manually curated four benchmark protein sets with known localizations from the literature, namely: Pyrenoid specific proteins (P; 18 proteins), proteins found in the pyrenoid and the stroma (PS; 10 proteins), proteins found in the stroma but excluded from the pyrenoid (S; 15 proteins), and non-chloroplast proteins (NC; 13 proteins) (Figure 2C and Supplemental Data Set 3). We used these benchmark proteins to calculate the enrichment threshold used to assess significant pyrenoid enrichment by applying a Receiver-Operator Characteristic (ROC) analysis ((Branon et al., 2018); Figure 2D). For the ROC analysis we adopted a stringent threshold by considering true positive proteins as exclusively pyrenoid localized (P) proteins. It should be noted that a portion of the pyrenoid localized proteins used for ROC analysis do not partition within the LLPS pyrenoid matrix but localize to the starch plate or the pyrenoid tubules. However, we reasoned that their close association to the pyrenoid would still support their labeling by RBCS2-TurboID.

**Figure 2.**
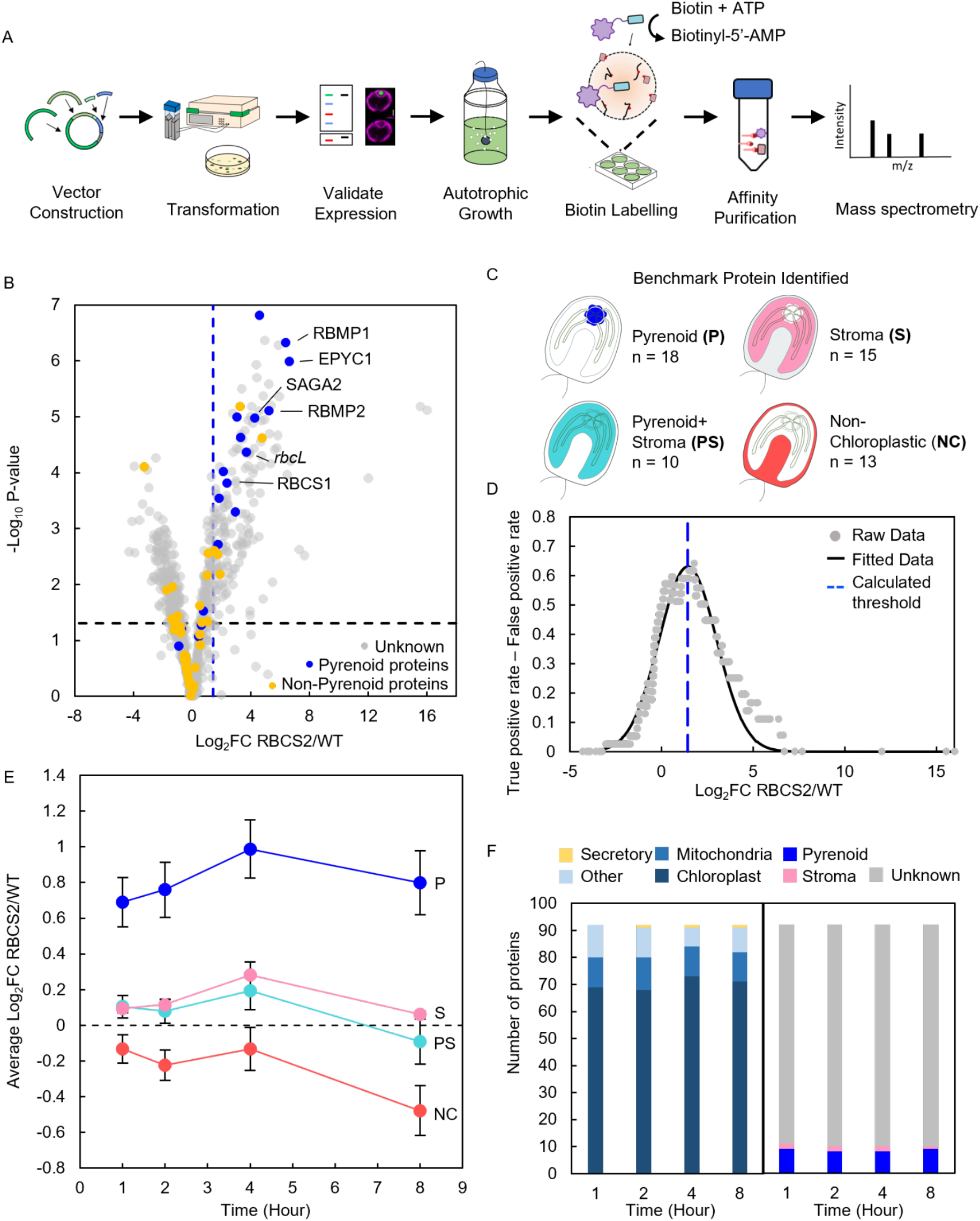
TurboID pipeline development and optimization of labeling time. **A**, Schematic representation of the developed TurboID pipeline. **B**, Volcano plot representing Log2 fold change (Log_2_ FC) between protein abundance in RBCS2-TurboID and WT. Proteins are colored according to their localization: unknown (gray), pyrenoid proteins (blue) and other localizations including chloroplast stroma, pyrenoid+stroma and non-chloroplastic (yellow). The Log_2_ FC threshold (dashed blue line) is calculated via the ROC analysis where only pyrenoid proteins are considered true positives. -Log_10_ P-value was used to represent statistical significance from the one-way Anova test carried out on the difference in abundance between RBCS2-TurboID and WT. P-value of <0.05 was used as a threshold. **C**, Benchmark proteins detected from the RBCS2-TurboID sample. **D**, Tradeoff between the true positive rate and false positive rates was plotted against the Log_2_ FC value, a gaussian function was fitted to the experimental data to determine a maximum, which is used as the enrichment threshold used in (B). **E**, Log_2_ FC of RBCS2-TurboID according to localization category in (C) was calculated at each labeling time point. **F**, PredAlgo predicted localization and benchmark protein categories of the top 10% enriched proteins from RBCS2-TurboID at each labeling time.

We then investigated protein labeling by RBCS2-TurboID at each time point for the different benchmark sets (Figure 2E). We saw that pyrenoid localized proteins (P) consistently showed the highest labeling across all time points. Chloroplast localized proteins (P, PS and S protein sets) exhibit an increase in labeling from 1 to 4 hours while non-chloroplastic proteins remain stable. Interestingly, all benchmark proteins appear to decrease in labeling at the 8 hour time point. This decrease is due to an increase in biotinylated protein abundance in untagged WT, rather than reduced labeling by RBCS2-TurboID (Supplemental Data Set 2).This data suggests that protein labeling in RBCS2-TurboID has approached full saturation within 4 hours. When we compared the top 10% of enriched proteins across the four timepoints, we found consistent enrichment for pyrenoid localized proteins, consistent overlap in protein identity >72% and agreement in their predicted cellular localization (Figure 2F; Supplemental Data Set 2).

Collectively, most pyrenoid localized proteins can be enriched within the first 2 hours, however increasing incubation time leads to increased biotinylation, with the largest differences between pyrenoid proteins and pyrenoid-excluded stromal proteins and non-chloroplast proteins occurring at 4 hours. We thus opted for 4 hour incubations for future experiments. We hypothesize that the rapid labeling dynamics of pyrenoid proteins within the first hour and the gradual increase in labeling of pyrenoid, pyrenoid+stroma and stroma can be explained by the LLPS properties of the pyrenoid where Rubisco is present in both the condensed phase (pyrenoid) and dilute phase (stroma). The high concentration of Rubisco in the condensed phase enables rapid labeling of proximal pyrenoid proteins. However, as Rubisco is also in the dilute phase at a much lower concentration, stromal proteins are gradually biotinylated over a longer time period.

### Stromal-TurboID controls enables a refined pyrenoid proteome

Although our current approach enabled enrichment of pyrenoid proteins, we set out to refine the pyrenoid proteome by trying to distinguish between pyrenoid specific proteins and proteins that are found within the pyrenoid and the stroma, and to remove the bias of increased labeling of abundant background proteins - a typical challenge in proximity labeling studies (Han et al., 2018). To achieve this we developed two chloroplast stromal controls and an additional pyrenoid specific TurboID line. For stromal controls we identified two Calvin-cycle enzymes, Ribulose Epimerase 1 (RPE1; Cre12.g511900) and Phosphoribulokinase 1 (PRK1, Cre12.g554800), that are abundant and localize to the chloroplast stroma but are excluded from the pyrenoid matrix (Figure 3A) (Küken et al., 2018). The Rubisco linker protein EPYC1 was chosen as an additional pyrenoid specific protein due to its abundance and functional importance for LLPS of Rubisco to form the pyrenoid (Mackinder et al., 2016). These new constructs; RPE1-TurboID, PRK1-TurboID, and EPYC1-TurboID; were assembled as C-terminal TurboID fusions and activity assessed in line with RBCS2-TurboID (Figure 3B and Supplemental Figure 2A and B).

**Figure 3.**
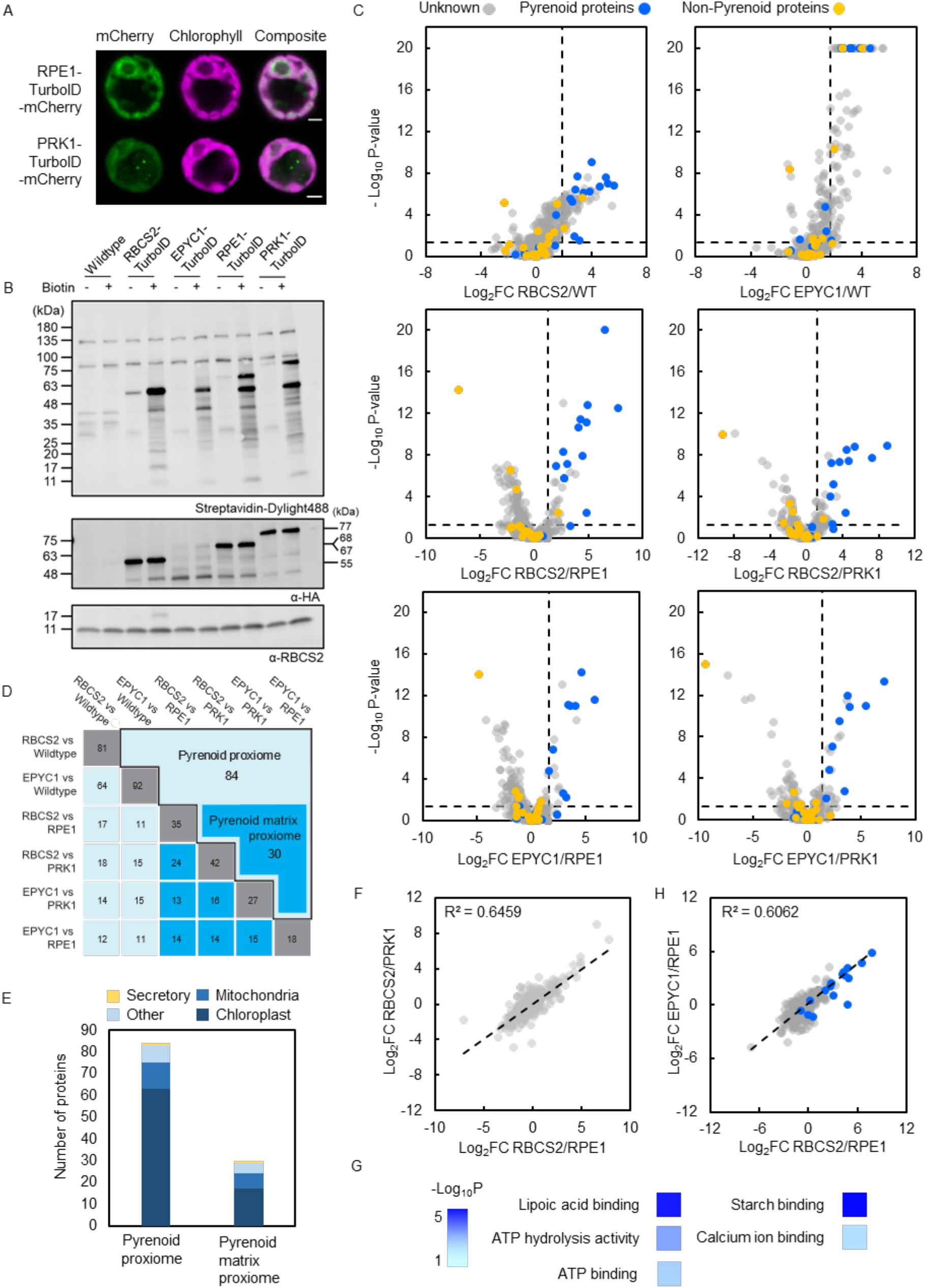
Determining the pyrenoid proteome using proximity labeling **A**, Localization of the mCherry fusion of RPE1- and PRK1-TurboID. Green and magenta signals represent the mCherry and chlorophyll autofluorescence respectively. Scale bar: 2 µm. **B**, Labeling activity of RBCS2-, EPYC1-, RPE1- and PRK1-TurboID lines were determined in the absence or presence of 2.5 mM biotin for 4 hours. Biotinylation was visualized via immunoblotting whole-cell lysate with streptavidin. Expression of RBCS2-TurboID (55 kDa), EPYC1-TurboID (68 kDa), RPE1-TurboID (67 kDa) and PRK1-TurboID (77 kDa) was probed by anti-HA. Anti-RBCS was used as a loading control. **C**, Volcano plots representing the Log_2_ Fold Change of RBCS2-TurboID and EPYC1-TurboID compared to WT and stromal controls. Pyrenoid proteins (blue dots) and non-pyrenoid proteins (yellow dots) were used to calculate the enrichment thresholds (vertical dashed line), the values are as follows: RBCS2/WT (1.88); RBCS2/RPE1 (1.31); RBCS2/PRK1 (1.14); EPYC1/WT (1.74); EPYC1/RPE1 (1.67); EPYC1/PRK1 (1.42). Statistical significance for each pairwise comparison was calculated using the PEAKSQ method, a significance P-value cut off of <0.05 was used (horizontal dashed line). The maximum -Log_10_ P-value computed by PEAKSQ is 20. **D**, Overlap matrix of identified proteins which are above the enrichment threshold in each treatment group. Bolded border highlights the overall pyrenoid proxiome while the dark blue shaded box denotes the pyrenoid matrix proxiome. For both the pyrenoid proxiome and pyrenoid matrix proxiome proteins had to be above the threshold in two or more comparisons. **E**, The distribution of predicted localization in both the pyrenoid proxiome and pyrenoid matrix proxiome obtained from PredAlgo (Tardif et al., 2012). **F**, Comparison of Log_2_ Fold Change in RBCS2-TurboID between the two stromal controls. **G**, Gene ontology (GO) enrichment analysis of the pyrenoid matrix proxiome (n=29) using the PANTHER GO Complete Molecular Function dataset. Significance as -Log_10_ P-value calculated from Fisher’ s exact test is presented in a color gradient. Only the GO Terms that were represented by two or more proteins are shown. **H**, Comparison of protein enrichment between RBCS2-TurboID and EPYC1-TurboID. Blue dots represent known pyrenoid proteins.

To ensure the optimal conditions for identifying the pyrenoid proteome, all expression lines were grown photoautotrophically in 0.04% CO2 where nearly all of the Rubisco is recruited to the pyrenoid and the CCM is fully induced (Mackinder, 2018). Labeling was allowed to proceed for 4 hours and resulting biotinylated proteins were enriched with streptavidin beads (see methods). Samples in triplicate were Tandem Mass Tag (TMT) labeled to enable a relative-quantification comparison of protein abundance between each line. We identified a total of 831 proteins derived from 5227 peptides with each protein containing at least 2 unique peptides. The Log2 Fold Change in reporter ion intensity was calculated between the pyrenoid-specific TurboID lines (RBCS2-TurboID and EPYC1-TurboID) and controls (WT, RPE1-TurboID and PRK1-TurboID). We then determined the enrichment of pyrenoid proteins in each comparison. In agreement with our previous pilot experiment, we found pyrenoid proteins are predominantly enriched by the pyrenoid-specific TurboID lines across all comparison groups (Figure 3C; blue dots).

To calculate the enrichment threshold used to assess significant pyrenoid enrichment, we applied the ROC analysis as in Figure 2C and D and a significance threshold of p-value <0.05 was calculated by the PEAKSQ significance test (Cox and Mann, 2008). We applied this analysis across all 6 of the comparison groupings (Figure 3C). In total this yielded 141 unique proteins across the 6 groups (Supplemental Data Set 5). To further filter out possible non-pyrenoid localized proteins, only identified proteins which are consistently above the enrichment threshold in at least 2 of the comparison groupings were considered true pyrenoid components. This leaves a final 84 unique proteins. We termed this the “pyrenoid proxiome” (Figure 3D, black bordered box). The pyrenoid proxiome contains 14 out of 19 known pyrenoid components detected in our dataset and is highly enriched for proteins that are predicted to be targeted to the chloroplast (Figure 3E).

We wondered whether comparison against stromal control lines improves distinction between pyrenoid proteins and stromal proteins relative to a WT control as hypothesized. We first tested if there were any major differences between our two stromal controls. Plotting the Log2 Fold Change of RBCS2-TurboID/RPE1-TurboID vs RBCS2-TurboID/PRK1-TurboID showed a strong correlation (R^2^=0.6459; Figure 3F), suggesting that both controls give similar results and that their similar stromal localization is the main driver of protein labeling. We next determined the difference between mean Log_2_ Fold Change of known pyrenoid and stromal proteins in each comparison pair (ie: RBCS2-TurboID vs WT or RBCS2-TurboID vs RPE1-TurboID). Indeed, the difference between mean Log_2_ fold change of pyrenoid and stromal proteins is most evident in the stromal control comparisons (Supplemental Figure 3). This is further supported by our observation that proteins peripheral to the pyrenoid Rubisco-EPYC1 matrix but not in it, such as LCIB, LCIC, STA2 and SBE3 (Yamano et al., 2010; Mackinder et al., 2017), are not enriched when stromal-specific TurboID lines are used as controls in place of WT. Our data here indicates that using the stromal controls gives a robust proteome of the Rubisco matrix. Taking proteins that are only seen above the threshold in two or more comparisons with stromal controls gives us 30 proteins (Supplemental Data Set 6). We named this the “pyrenoid matrix proxiome” (Figure 3D).

GO term enrichment analysis of the pyrenoid matrix proxiome indicated that proteins could be functionally grouped into a small number of processes (Figure 3G). This includes Lipoic acid binding which represents sulfur related compounds (GO: GO:0031405), carbohydrate related processes like Alpha-amylase activity and starch binding (GO: GO:0004556 and GO:2001070 respectively) and ATP-binding groups (GO:0005524). We found multiple proteins in the pyrenoid matrix proteome, notably Cre06.g269650, Cre03.g158050 and SAGA1 that contain a starch binding domain alongside a variety of functional domains. This argues that the matrix-starch interface might act as a specialized site for specific structural or biological processes. A broader analysis of the pyrenoid proxiome also reveals that multiple proteins contain iron-sulfur binding domains (Cre05.g240850, Cre13.g592200 and Cre02.g093650) and have RNA-related functions (Cre10.g440050, Cre10.g435800, Cre09.g393358, Cre13.g578650). Tentatively, enrichment of these proteins in the pyrenoid proxiome suggests that the pyrenoid might take on additional roles to carbon fixation.

### RBCS-TurboID and EPYC1-TurboID result in comparable pyrenoid proteomes

Rubisco and EPYC1 are the two core proteins of the pyrenoid, interact with each other and are both essential for phase separation and pyrenoid formation. However, an APMS study using both RBCS2 (and RBCS1) and EPYC1 as baits identified multiple distinct interacting partners as well as a shared set of interactors (Mackinder et al., 2017). Since the majority of pyrenoid proteins we have used as a benchmark in this study were previously characterized due to their interactions with the RBCS, it is difficult to ascertain whether the use of RBCS2-TurboID has preferentially labeled Rubisco interactors or the more broader pyrenoid proteome. We reasoned that by comparing proteins obtained from RBCS2-TurboID against that of EPYC1-TurboID we might be able to provide the distinction and more broadly determine if proximity labeling of proteins in a dynamic molecular condensate preferentially labels the proteome of the condensate vs direct interactors of the bait. A comparison of EPYC1-TurboID and RBCS2-TurboID’ s respective fold change against the stromal controls showed a strong correlation (R^2^ = 0.6062), this was considerably strengthened when we focus on known pyrenoid proteins (R^2^ = 0.7054; Figure 3H, blue dots). In conclusion, irrespective of bait, using a mobile protein of the phase separated pyrenoid gives a high-confidence proteome of a biomolecular condensate.

### Proximity labeling identified novel pyrenoid proteins

To validate our pyrenoid proteome, 8 previously unlocalized proteins were selected from preliminary data and our initial RBCS2-TurboID vs WT comparison for fluorescence tagging (Figure 4A). Target genes were cloned in frame with a Venus or mScarlet-I fluorescence protein by recombineering that retains their native promoter (Emrich-Mills et al., 2021), then transformed into WT *Chlamydomonas*. Seven of the tagged proteins showed a pyrenoid only localization with a broad range of sub-pyrenoid localization patterns shown (Figure 4B).

**Figure 4.**
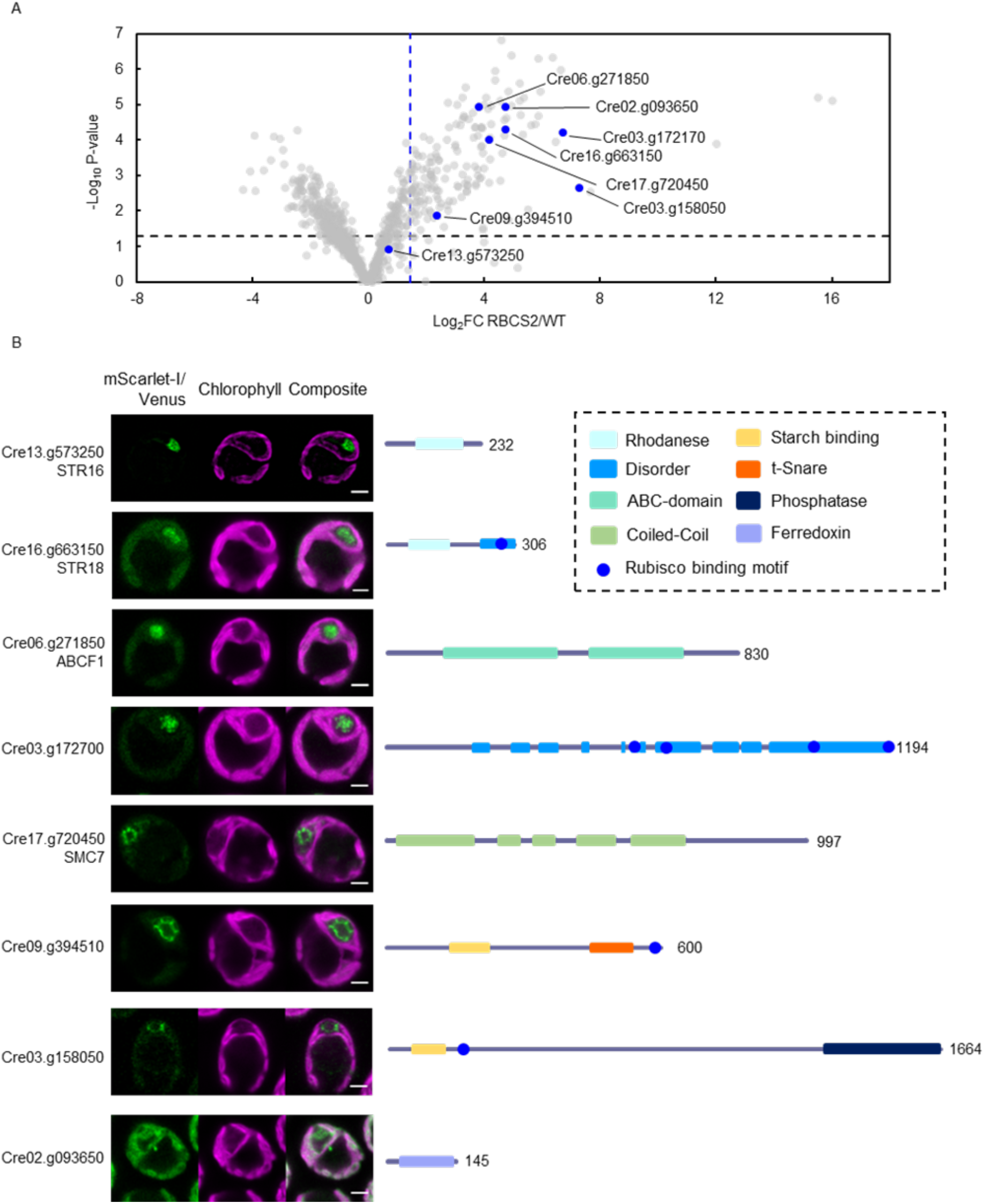
Proximity labeling identifies novel pyrenoid proteins. **A**, The Volcano plot in Figure 2B was reproduced here to highlight the proteins that were chosen for localization (blue dots). **B**, Confocal imaging of the chosen proteins. Proteins were expressed as Venus or mScarlet-I fusions using their native promoter sequence. Green and magenta signals denote the fluorescence channel and autofluorescence from chlorophyll. Scale bar: 2 µm. Schematic overview of structural prediction from PSI-pred and conserved domains are highlighted next to the confocal images.

STR16 (Cre13.g573250), STR18 (Cre16.g663150) and ABCF1 (Cre06.g271850) showed a localization pattern consistent to the pyrenoid matrix, which is supported by their lack of a predicted transmembrane or starch binding domain. STR16 and STR18 contain a catalytically active rhodanese (thiosulfate sulfurtransferase) domain as opposed to CAS1 and RBMP2 that have previously been localized to pyrenoid but lack a critical cysteine in their Rhodanese domains and are thus presumably catalytically inactive. Rhodanese domains have been implicated in an array of functions including disulfide bond formation (Chng et al., 2012) and iron-sulfur (Fe-S) cluster biosynthesis (Bonomi et al., 1977). The latter is particularly interesting as multiple proteins in the pyrenoid proxiome contain an iron-sulfur cluster domain (Figure 3G). ABCF1 (Cre06.g271850) is predicted to be an ATP binding cassette family F like protein (ABCF) which has been shown to regulate protein translation via their binding to ribosomes (Boël et al., 2014). The ABCF annotation is also supported by AlphaFold modeling (Supplemental Figure 4, (Jumper et al., 2021)), which suggests ABCF1 contains the canonical arm and linker domain. Cre03.g172700 forms distinct puncta within the pyrenoid matrix unlike the more homogenous signal observed for matrix proteins such as RBCS2. This sub-pyrenoid localization suggests that it may be associated with the pyrenoid tubules. While PSI-pred structural prediction suggests Cre03.g172700 is predominantly disordered, AlphaFold prediction suggests that its latter half is composed of a central long alpha-helix surrounded by multiple shorter helices interspaced with disordered sequences that in total contain 4 RBMs (Supplemental Figure 4). Unlike the other proteins which localize to the pyrenoid matrix, SMC7 (Cre17.g720450), Cre09.g394510 and Cre03.g158050 are found at the edge of the pyrenoid matrix, with SMC7 and Cre03.g158050 forming discrete puncta surrounding the matrix while Cre09.g394510 appears to line the starch-matrix interface. These proteins show a similar localization pattern to SAGA1 which occupies the starch-matrix-tubule interface. SMC7 is predicted to be a structural maintenance of chromosomes (SMC) protein. However, SMC7 lacks the signature ATP-binding and hinge domain important for its function in chromatin condensation (Harvey et al., 2002), and instead only contains the conserved coiled-coil domain. This structure arrangement mirrors that of SAGA1 and SAGA2 (Itakura et al., 2019) that were also annotated with a SMC prediction. Both Cre09.g394510 and Cre03.g158050 contain a CBM20 starch binding domain. Cre09.g394510 possess a t-Snare domain at its C-terminal, which is known to mediate vesicle fusion processes (Han et al., 2017), suggesting that Cre09.g394510 could be involved in membrane remodeling of the pyrenoid tubules as they are structurally reorganized from thylakoid sheets to pyrenoid tubules as they traverse gaps within the starch sheath (Engel et al., 2015). On the other hand, the C-terminal region of Cre03.g158050 contains a protein phosphatase domain (PPM-type phosphatase). While the phosphorylation of EPYC1 in low CO2 conditions has been proposed to regulate binding to Rubisco (Turkina et al., 2006; Barrett et al., 2021), such phosphorylation is not necessary for EPYC1 to phase separate Rubisco (Wunder et al., 2018). The localization of Cre03.g158050 specifically to the pyrenoid periphery suggests that there may be a sub-pyrenoid spatial control of pyrenoid protein phosphorylation state.

Together, the discovery of novel pyrenoid components with diverse localization and functional annotations provides us with additional candidates which could have a role in pyrenoid formation.

### Changes in the pyrenoid proteome in response to CO_2_

At high CO_2_, when a CCM is not required the pyrenoid partially dissolves, with ∼50% of Rubisco leaving the pyrenoid into the surrounding stroma (Borkhsenious et al., 1998). In addition, the starch sheath breaks down and stromal starch increases (Kuchitsu et al., 1988). However, at a transcriptional and protein abundance level, matrix pyrenoid proteins show a broad range of responses (Fang et al., 2012; Brueggeman et al., 2012; Arias et al., 2020). To explore if the pyrenoid composition changes in response to CO_2_ we compared RBCS2-TurboID lines grown at both high and low CO_2_ (Figure 5A and B; Supplemental Data Set 7). A large number of previously known pyrenoid proteins and proteins in our pyrenoid matrix proxiome were not preferentially enriched indicating that the vast majority of the pyrenoid proteome is not CO_2_ responsive. However, a small number of proteins showed a greater than 2-fold change, with 20.5% (7/34) enriched at low CO_2_ and 2.9% (1/34) enriched at high CO_2_. Three of the low CO_2_ enriched proteins, SAGA1, LCI9 and AMA3, are associated with starch binding/metabolism. LCI9 has previously been localized to the starch plate interfaces and proposed to play a role in starch breakdown (Mackinder et al., 2017). AMA3 is an alpha amylase also involved in starch hydrolysis (Gargouri et al., 2015) and mutants in SAGA1 have a severe starch structural defect (Itakura et al., 2019). Collectively, this supports the major remodeling of starch to form the starch sheath under low CO_2_ conditions.

**Figure 5.**
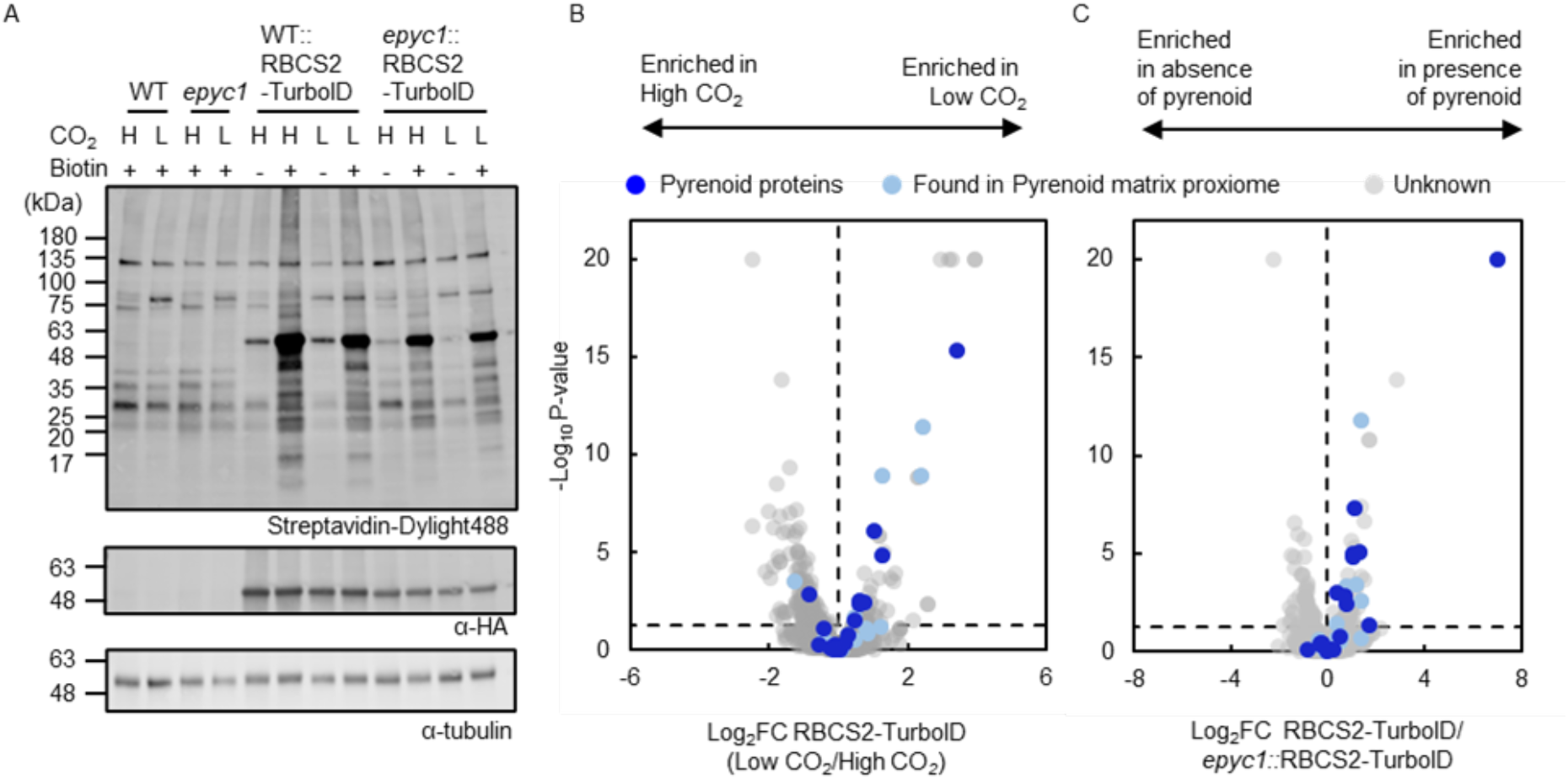
Proximity labeling suggests the pyrenoid proteome has a subtle response to changes in CO_2_ and phase separation. **A**, Protein labeling of RBCS2-TurboID expressed in WT and *epyc1* as well as their corresponding untagged background were tested under different CO_2_ conditions. Respective cell lines were grown photoautotrophically and supplemented with 3% CO_2_ (H) or 0.04% CO_2_ (L). Harvested cells were incubated with 2.5 mM biotin for 4 hours. Labeling was visualized by immunoblotting the whole cell lysate against streptavidin. Anti-HA was used to probe for RBCS2-TurboID expression and anti-tubulin was used as a loading control. **B-C**, Volcano plots representing the Log_2_ Fold Change of RBCS2-TurboID in low CO_2_ vs high CO_2_ (**B**); or RBCS2-TurboID expressed in WT background compared to RBCS2-TurboID expressed in the *epyc1* mutant **(C)**. Known pyrenoid proteins and the pyrenoid matrix proxiome are colored dark blue and light blue respectively, while unknowns are colored in gray. Statistical significance for each pairwise comparison was calculated using the PEAKSQ method, a significance cut off for P-value <0.05 was used (horizontal dashed line).

### Possible role of phase separation in recruitment to the pyrenoid matrix

The deletion of EPYC1 leads to abolishment of the pyrenoid and CCM due to the failure to condense Rubisco into the pyrenoid (Mackinder et al., 2016). Confident that the pyrenoid matrix proxiome is labeled by RBCS2-TurboID we explored how labeling changed upon condensation of Rubisco into the pyrenoid. To achieve this we compared RBCS2-TurboID expressed to similar levels in both WT and the *epyc1* mutant (Figure 5A). A large number of known pyrenoid proteins and pyrenoid matrix proxiome proteins are enriched in WT vs *epyc1* (Figure 5C; Supplemental Data Set 7), indicating that phase separation either results in more efficient labeling or that phase separation is required for close proximity to Rubisco. A subset of proteins showed very little enrichment upon Rubisco condensation, suggesting that these proteins may directly interact with Rubisco independent of pyrenoid formation. Unexpectedly, many Rubisco binding motif containing proteins require pyrenoid formation to be enriched (top right quadrant of Figure 5C) indicating that the weak binding affinity (Kd ∼3 mM; (He et al., 2020)) of RBMs may not be sufficient to allow Rubisco-RBM complex formation prior to Rubisco condensation by EPYC1.

## Discussion

We have established TurboID-based proximity labeling in the chloroplast of the model green alga *Chlamydomonas reinhardtii*. Proximity labeling has proven powerful in unraveling a broad range of cellular processes and suborganelle composition in a diverse range of organisms including plants (Mair and Bergmann, 2022; Zhang et al., 2019), diatoms (Turnšek et al., 2021) and cyanobacteria (Dahlgren et al., 2021). However, until now it has not been established in plastids or *Chlamydomonas*. The parallel manuscripts in this issue give a snapshot of the diversity of possible applications of TurboID in both plant (Wurzinger et al.) and algal plastids (our study and Kreis et al.). The independently-determined similar biotin concentrations and incubation time for labeling in the *Chlamydomonas* chloroplast by our work and the work by Kreis et al. highlight the reproducibility and robustness of the method.

Once established we applied TurboID to determine the protein composition of the phase-separated pyrenoid. Our data has identified a “pyrenoid proxiome” containing 84 proteins. A large number of previously localized pyrenoid proteins (67%) from the literature are seen in our pyrenoid proxiome. However, it does miss several previously classified pyrenoid proteins. A deeper analysis of these proteins indicate that they are within specific pyrenoid sub-domains that matrix generated biotin radicals potentially fail to penetrate due to either spatial or physical constraints. For example, CAH3, a pyrenoid tubule lumen protein is not identified in our proxiome most likely due to the limited penetration of biotin radicals across membranes (Rhee et al., 2013).

By including robust stromal controls of proteins that are adjacent to the pyrenoid but do not partition into the matrix we could determine the “pyrenoid matrix proxiome” containing 30 proteins. This protein set excluded multiple proteins classed as pyrenoid proteins that are found at the pyrenoid periphery but do not partition into the matrix. These include LCIB, LCI9, LCIC, SBE3. This data along with the identification of nearly all known matrix proteins and proteins with RBMs that are at the matrix interface (i.e. SAGA1, BST4, RBMP2) gives us high confidence in this dataset.

GO-term analysis of the pyrenoid matrix proxiome and a broader analysis of the pyrenoid proxiome shows the enrichment of proteins in a small number of biochemical processes suggesting that the pyrenoid plays additional roles to CO_2_ concentration. Three groups that stood out were RNA binding/translation proteins, Fe-S containing proteins and starch binding proteins. Biomolecular condensates are regularly associated with RNA sequestration and processing (Banani et al., 2017). This association allows cells to respond in a timely manner in face of cellular stress. In *Chlamydomonas* photosynthetic machinery is translated at a specialized position adjacent to the pyrenoid called the T-zone (or Translation zone; (Sun et al., 2019)). Under light and oxidative stress, the core photosystem II component PsbA mRNA becomes enriched within the pyrenoid matrix (Uniacke and Zerges, 2008; Zhan et al., 2015) which suggests the pyrenoid recruits RNA as a stress response. However, the molecular basis and function of this mRNA sequestration remains unclear. In this study, multiple RNA-associated proteins are found within the pyrenoid proxiome (Cre10.g440050, Cre10.g435800, Cre09.g393358, Cre13.g578650). We have also localized a novel ribosome-associated protein ABCF1 to the pyrenoid. An *E*.*coli* homologue of ABCF1, EttA, was demonstrated to prevent translation by its binding to 70s ribosomes in a ATP/ADP ratio dependent manner (Boël et al., 2014). The localization of ABCF1 to the pyrenoid further supports a role of the pyrenoid in RNA metabolism where it could either act to sequester chloroplastic ribosomes in the pyrenoid or partition ABCF1 away from chloroplastic ribosomes under certain environmental conditions.

Fe-S protein assembly and activity is typically sensitive to molecular O_2_ (Boyd et al., 2014). It was intriguing to see that the pyrenoid was enriched for both Fe-S assembly and Fe-S containing proteins. A proposed, but unconfirmed, function of the pyrenoid to enhance CO_2_ fixation is to minimize the presence of O_2_ to increase the CO_2_:O_2_ ratio at the active site of Rubisco. A reduced O_2_ environment could also favor other O_2_ sensitive biological processes. We found that the rhodanese domain containing proteins STR16 and STR18 were localized into the pyrenoid, with rhodanese domains linked to the biogenesis of Fe-S clusters (Rydz et al., 2021). Pyrenoid localization might allow them to be shielded from the oxygenic environment outside the pyrenoid matrix, allowing these oxygen sensitive processes to be carried out. Alternatively, rhodanese also has been suggested to participate in reactive oxygen species (ROS) scavenging via the production of reactive sulfur species (Wang et al., 2021). Since ROS has also been found to drive pyrenoid formation (Neofotis et al., 2021), the presence of rhodanese domain containing proteins in the pyrenoid suggests that the pyrenoid itself is involved with ROS metabolism or redox signaling.

The pyrenoid starch sheath is proposed to act as a diffusion barrier which limits CO_2_ diffusion away from the pyrenoid matrix. Recent evidence has suggested that this matrix- starh association is critical for the organization of many pyrenoid components. The deletion of a starch-binding protein SAGA1 results in the formation of multiple pyrenoids with altered starch sheath and pyrenoid tubule morphology (Itakura et al., 2019). And the knock-out of Isoamylase 1 (ISA1) that abolishes the pyrenoid starch sheath results in the CCM-essential carbonic anhydrase LCIB to mis-localize as an aggregate at the basal region of pyrenoid (Toyokawa et al., 2020), in contrast to its typical pyrenoid periphery localization. Together, starch binding proteins show crucial importance to the functioning of the pyrenoid in CCM-related processes. In this work, two additional proteins (Cre09.g394510, Cre03.g158050) that contain a starch binding CBM20 domain have been localized to the pyrenoid. Both proteins possess a second functional domain, with this domain arrangement also seen in SAGA1 and LCI9 which incidentally share a similar localization pattern (Mackinder et al., 2017). Investigating the role of these proteins in pyrenoid structural organization and function could provide novel insights into pyrenoid assembly needed for future engineering of a functional pyrenoid into higher plants (Adler et al., 2022).

Once we had determined a high-confidence pyrenoid proteome we explored the change in the proxiome of Rubisco at both low and high CO_2_ and with and without phase separation. Surprisingly most proteins appear to be present in the pyrenoid under both low and high CO_2_ conditions indicating that the core proteome of the pyrenoid is relatively stable. However, a subset involved in starch metabolism are predominantly enriched under low CO_2_ when starch needs to be remodeled to form a CO_2_ leakage barrier. By using the *epyc1* mutant we explored how labeling by RBCS2-TurboID changed upon Rubisco condensation into the pyrenoid. Most pyrenoid matrix proxiome components were enriched by Rubisco condensation indicating that they are brought into closer proximity upon pyrenoid formation. A subset however showed very little change suggesting that they may already be interacting with Rubisco independent of pyrenoid assembly. For both the high vs low CO_2_ and WT vs *epyc1* comparisons it should be noted that the 4 hour incubation time of labeled strains could have led to translational changes resulting in compounding data between absolute protein amount and partitioning into the pyrenoid. In addition, the partial dissolution of the pyrenoid during high CO_2_ also results in a higher proportion of RBCS2-TurboID in the dilute phase. This in turn potentially increases labeling of proteins which have yet partitioned into the pyrenoid. In the future shorter labeling times could help further refine the pyrenoid proteome under varying conditions.

Proximity labeling has been underutilized for understanding phase separated proteomes that are highly dynamic and thus are challenging to purify (Hubstenberger et al., 2017). The presence and exchange of bait proteins between the condensed phase and dilute phase might result in reduced specificity of RBCS2/EPYC1 -TurboID over time and labeling outside of the condensate. To counteract this, we found that the use of abundant soluble controls that are excluded from the pyrenoid enabled a highly refined pyrenoid matrix proteome to be determined. Future experiments using proximity labeling, specifically to determine the proteomes of biomolecular condensates, should include carefully chosen controls.

To make TurboID easily accessible for other labs using *Chlamydomonas* we based our cloning on the MoClo golden gate cloning framework that enables TurboID to be used with a broad range of parts (Crozet et al., 2018) and easily fused to proteins that are already within this framework. To enable easy adoption of this powerful method all developed vectors and lines are deposited at the *Chlamydomonas* Resource Centre.

## Methods

### Construction of APEX2/TurboID expression vector in Chlamydomonas

Construction of APEX2/TurboID-expression cassettes in *Chlamydomonas* is designed using the modular cloning system *Chlamydomonas* MoClo toolkit (Crozet et al., 2018). Golden Gate compatible syntaxes were added to synthesized parts encoding the APEX2/TurboID enzyme and target genes (RBCS2/EPYC1), or via PCR using CC-4533 genomic DNA for RPE1/PRK1. Due to the low complexity and high repeat nature of EPYC1, the EPYC1 coding sequence was synthesized in 4 parts as a Level -1 construct, while RBCS2 was synthesized as two parts to avoid a detected sequence repeat. The APEX2 and TurboID tags (Branon et al., 2018; Ganapathy et al., 2018) were codon optimized for *Chlamydomonas* (Nakamura et al., 2000) with the RBCS2i2 (Cre02.g120150) and LHCBM1i2 (Cre01.g066917) introns inserted at approximately 500 bp increments to improve protein expression (Baier et al., 2018). The enzyme tags were similarly synthesized as Level -1 parts. Together, the Level -1 and PCR amplified target gene, APEX2/TurboID tag and a small flexible linker (GSGSTSGSGS) were assembled to a Level 0 product occupying the B3-B4 MoClo position using the pUAP1 backbone such that the target genes are expressed with the enzyme tag at its C-terminal that is bridged by a small flexible linker. The Level 1 cassette was then assembled using the target gene-TurboID/APEX2 fusion part, the PSAD Promoter/Terminator pair and either a C-terminal tandem HA/Flag tag epitope for labeling experiments or a mCherry tag for localization. The resultant Level 2 expression module consists of target gene-TurboID fusion cassette and an antibiotic resistance cassette for selection. To enable accessible use of TurboID-based proximity labeling in the golden gate cloning pipeline, the identical TurboID coding sequence with the flexible linker was also cloned into a Level 0 part occupying the B4 MoClo position. Sequences for all developed vectors are in Supplemental Data Set 1. All vectors and strains will be deposited at the Chlamydomonas Resource Centre prior to publication.

### Chlamydomonas growth and transformation

*Chlamydomonas* cultures were maintained on Tris-phosphate acetate (TAP) medium with revised Hunter’ s trace element (Kropat et al., 2011) unless mentioned otherwise. Assembled plasmids were linearised with I-SceI (for fluorescent tagging plasmids) or BsaI (for proximity labeling plasmids) and transformed into *Chlamydomonas* via electroporation according to (Mackinder et al., 2017).

### Protein extraction and Immunoblotting

For immunoblotting, photoautotrophically grown cells at mid-log phase were harvested by centrifugation x17,900g for 5 minutes at 4°C. Cell pellets were resuspended in lysis buffer (25 mM Tris-HCl pH7.4, 500 mM NaCl, 1 mM DTT, 5 mM MgCl2, 0.1 mM PMSF, 1x EDTA-free protease inhibitor (Roche), 0.1% SDS, 0.5% Deoxycholic Acid, 1% Triton-X100) before snap-freezing in liquid nitrogen. The cell suspensions were lysed by 5 freeze/thaw cycles. Supernatants were used as protein extract after centrifugation x17,900g for 10 minutes at 4°C and then stored at -70°C. For immunoblotting, boiled protein samples were resolved by SDS-PAGE and transferred to a PVDF membrane via a semi-dry transfer. Membrane was blocked with 3% BSA in Tris Buffered Saline with 0.1% Tween 20 (TBST) and probed with antibodies accordingly. Antibodies are diluted in TBST as follows: Streptavidin Dylight-488 conjugate (1:4000, Fisher scientific #21832); anti-HA (1:1000, Fisher scientific 26183); anti-Flag (1:1000, Sigma #F1804); anti-Tubulin (1:2000, Sigma #T6074).

### Biotin Labeling and Streptavidin-affinity purification

All three TurboID-labeling experiments were performed similarly. The starter culture of TurboID expression lines and wildtype were grown to mid-log phase in TAP medium. They were used to inoculate 400 mL of Tris-Phosphate (TP) medium that was supplied with air-level CO_2_ (0.04% CO_2_) or elevated CO_2_ (3% CO_2_) if indicated. Cells were harvested by centrifugation x1,500g for 5 minutes at room temperature. They were then resuspended in fresh TP medium in a 6-well cell culture plate to an OD_750_ of 2.5. 100 mM Biotin stock in DMSO was added to the cell suspension to a final concentration of 2.5 mM to initiate the labeling reaction. The biotin labeling was allowed to proceed for 1-8 hours in the pilot experiment or for 4 hours in the later experiments on an orbital shaker. Biotin labeled cells were harvested by centrifugation x21,300g, 2 minutes at 4°C after rinsing 3 times with ice-cold TP medium. Cell pellets were snap-frozen in liquid nitrogen and stored at -70°C until Streptavidin affinity purification.

For APEX2 labeling, the RBCS2-APEX2 expression cells were grown and harvested to an OD_750_ of 2.5 as mentioned above. Biotin-phenol at a final concentration of 2.5 mM was added to the harvested cell suspension from a 250 mM Biotin-Phenol stock in DMSO. Biotin-Phenol incubation was performed for 2 hours on an orbital shaker. The H_2_O_2_ activator at 2 mM concentration was spiked into the suspension to initiate the biotin labeling for 2 minutes. Reaction was then quenched by resuspending cell suspension in an ice cold quencher solution (10 mM sodium ascorbate, 5 mM Trolox and 10 mM sodium azide in PBS, pH 7.4) and pelleted by centrifugation x21,300g for 1 minute at 4°C and stored at -70°C until Streptavidin affinity purification.

Protein extraction was carried out as described above. Prior to streptavidin affinity pulldown free biotin was removed from protein samples using a Zeba™ Spin Desalting column (#89891, Thermofisher). To determine their protein concentration, a small aliquot (50 µL) of the desalted proteins were diluted by 10 fold in water and their abundance was measured via Pierce BCA protein assay kit (#23225, Thermofisher) as per manufacturer’ s instructions. For Streptavidin affinity purification, a total of 1.75 mg of protein was used with 50 uL of Pierce™ Streptavidin Magnetic Beads (88816; Thermofisher) equilibrated with lysis buffer. The bead suspension was incubated at 4°C overnight on a rotor wheel. Beads were then washed twice with lysis buffer for 5 minutes; once with 1 M KCl for 2 minutes; once with 0.1 M NaCO3 for 1 minute; once with 4 M Urea in 50 mM TEAB (Triethylammonium bicarbonate, pH 8.5) for 1 minute; once with 6 M Urea in 50 mM TEAB for 1 minute and twice with 50 mM TEAB buffer for 5 minutes. Washed beads were frozen at -70°C until submitted for mass spectrometry.

### LC-MS/MS and analysis of APEX2 and TurboID pilot studies

#### APEX2 digestion

For the APEX2 experiments, Streptavidin beads were eluted by boiling with a 2x Laemmli loading buffer (Biorad, 161-0737) supplemented with 20 mM DTT and 2 mM Biotin. The eluate was then ran on a 4-15% Tris-Glycine gel (Biorad, #4561084) for 30 minutes on 50V. Gel slices were then fixed according to (Mackinder et al., 2017). In-gel tryptic digestion was performed after reduction with 10 mM dithioerythritol and 50 mM *S*-carbamidomethylation with iodoacetamide. Gel pieces were washed two times with aqueous 50% (v:v) acetonitrile containing 25 mM ammonium bicarbonate, then once with acetonitrile and dried in a vacuum concentrator for 20 min. A 500 ng aliquot of sequencing-grade trypsin (Promega) was added prior to incubation at 37°C for 16 h.

#### TurboID Digestion

For the TurboID pilot experiment, on-bead digestion was performed after reduction with 10 mM tris(2-carboxyethyl)phosphine and alkylation with 10mM Iodoacetamide in 50 mM triethylamonium bicarbonate TEAB containing 0.01% ProteaseMAX surfactant (Promega). A 500 ng aliquot of sequencing-grade trypsin (Promega) was added prior to incubation at 37°C for 16 h.

#### LC-MS Acquisition APEX2 and TurboID

Resulting peptides were re-suspended in aqueous 0.1% trifluoroacetic acid (v/v) then loaded onto an mClass nanoflow UPLC system (Waters) equipped with a nanoEaze M/Z Symmetry 100 Å C_18_, 5 µm trap column (180 µm x 20 mm, Waters) and a PepMap, 2 µm, 100 Å, C_18_ EasyNano nanocapillary column (75 mm x 500 mm, Thermo). The trap wash solvent was aqueous 0.05% (v:v) trifluoroacetic acid and the trapping flow rate was 15 µL/min. The trap was washed for 5 min before switching flow to the capillary column. Separation used gradient elution of two solvents: solvent A, aqueous 0.1% (v:v) formic acid; solvent B, acetonitrile containing 0.1% (v:v) formic acid. The flow rate for the capillary column was 300 nL/min and the column temperature was 40°C. The linear multi-step gradient profile was: 3-10% B over 7 mins, 10-35% B over 80 mins, 35-99% B over 10 mins and then proceeded to wash with 99% solvent B for 8 min. The column was returned to initial conditions and re-equilibrated for 15 min before subsequent injections. The nanoLC system was interfaced with an Orbitrap Fusion Tribrid mass spectrometer (Thermo) with an EasyNano ionization source (Thermo). Positive ESI-MS and MS^2^ spectra were acquired using Xcalibur software (version 4.0, Thermo). Instrument source settings were: ion spray voltage, 1,900 V; sweep gas, 0 Arb; ion transfer tube temperature; 275°C. MS^1^ spectra were acquired in the Orbitrap with: 120,000 resolution, scan range: *m/z* 375-1,500; AGC target, 4e^5^; max fill time, 100 ms. Data dependent acquisition was performed in top speed mode using a 1 s cycle, selecting the most intense precursors with charge states >1. Easy-IC was used for internal calibration. Dynamic exclusion was performed for 50 s post precursor selection and a minimum threshold for fragmentation was set at 5e^3^. MS^2^ spectra were acquired in the linear ion trap with: scan rate, turbo; quadrupole isolation, 1.6 *m/z*; activation type, HCD; activation energy: 32%; AGC target, 5e^3^; first mass, 110 *m/z*; max fill time, 100 ms. Acquisitions were arranged by Xcalibur to inject ions for all available parallelizable time.

#### Spectral Counting APEX2

Peak lists in Thermo .raw format were converted to .mgf using MSConvert (version 3.0, ProteoWizard) before submitting to database searching against 19,716 *Chlamydomonas* protein sequences appended with common proteomic contaminants. Mascot Daemon (version 2.6.0, Matrix Science) was used to submit the search to a locally-running copy of the Mascot program (Matrix Science Ltd., version 2.7.0). Mascot was searched with a fragment ion mass tolerance of 0.50 Da and a parent ion tolerance of 3.0 PPM. O-124 of pyrrolysine, j-16 of leucine/isoleucine indecision and carbamidomethyl of cysteine were specified in Mascot as fixed modifications. Oxidation of methionine was specified in Mascot as a variable modification. Scaffold (version Scaffold_5.2.0, Proteome Software Inc., Portland, OR) was used to validate MS/MS based peptide and protein identifications. Peptide identifications were accepted if they could be established at greater than 84.0% probability to achieve an FDR less than 1.0% by the Percolator posterior error probability calculation. Protein identifications were accepted if they could be established at greater than 6.0% probability to achieve an FDR less than 1.0% and contained at least 2 identified peptides. Quantitative value of total spectra was used to calculate the Log_2_ Fold change between RBCS2-APEX2 and WT samples, and the Student’ s T-test derived P-value was -log_10_ transformed before presented.

#### Precursor Intensity-based Relative Quantification TurboID pilot

Peak lists in .raw format were imported into Progenesis QI (Version 2.2., Waters) and LC-MS runs aligned to the common sample pool. Precursor ion intensities were normalised against total intensity for each acquisition. A combined peak list was exported in .mgf format for database searching against 19,716 *Chlamydomonas* protein sequences appended with common proteomic contaminants. Mascot Daemon (version 2.6.0, Matrix Science) was used to submit the search to a locally-running copy of the Mascot program (Matrix Science Ltd., version 2.7.0). Search criteria specified: Enzyme, trypsin; Max missed cleavages, 1; Fixed modifications, Carbamidomethyl (C); Variable modifications, Oxidation (M),; Peptide tolerance, 3 ppm; MS/MS tolerance, 0.5 Da; Instrument, ESI-TRAP. Peptide identifications were passed through the percolator algorithm to achieve a 1% false discovery rate assessed against a reverse database and individual matches filtered to require minimum expect score of 0.05. The Mascot .XML result file was imported into Progenesis QI and peptide identifications associated with precursor peak areas and matched between runs. Relative protein abundance was calculated using precursor ion areas from non-conflicting unique peptides. Accepted protein quantifications were set to require a minimum of two unique peptide sequences. Missing values were then replaced by the minimal value detected from each bait. The fold change in RBCS2-TurboID vs WT comparison was calculated on the sum of relative protein abundance at all time points and were Log2 transformed. Statistical testing was performed in Progenesis QI from ArcSinh normalized peptide abundances and the ANOVA-derived p-values was -log10 transformed and presented.

### LC-MS/MS and analysis of TMT labeled TurboID experiments

#### TurboID digestion and TMT labeling

For the TurboID experiments in Figure 3 and 4, on-bead digestion was performed after reduction with 10 mM tris(2-carboxyethyl)phosphine and alkylation with 50 mM methyl methanethiosulfonatein in 50 mM triethylamonium bicarbonate TEAB. A 500 ng aliquot of sequencing-grade trypsin (Promega) was added prior to incubation at 37°C for 16 h. Post digestion the peptide containing supernatants were removed from the beads for TMT labeling. Peptides were labeled with TMTPro 16-plex reagents (Thermo) as detailed in the manufacturer’ s protocol. Post labeling samples were combined and dried in a vacuum concentrator before reconstituting in 100 ml H_2_O.

#### LC-MS Acquisition of TMT labeled TurboID experiment

Peptides were fractionated by high pH reversed phase C_18_ HPLC. Samples were loaded onto an Agilent 1260 II HPLC system equipped with a Waters XBridge 3.5 µm, C_18_ column (2.1 mm x 150 mm, Thermo). Separation used gradient elution of two solvents: solvent A, aqueous 0.1% (v:v) ammonium hydroxide; solvent B, acetonitrile containing 0.1% (v:v) ammonium hydroxide. The flow rate for the capillary column was 200 mL/min and the column temperature was 40°C. The linear multi-step gradient profile for the elution was: 5-35% B over 20 mins, 35-80% B over 5 mins, the gradient was followed by washing with 80% solvent B for 5 min before returning to initial conditions and re-equilibrating for 7 min prior to subsequent injections. Eluant was collected at 1 min intervals into LoBind Eppendorf tubes. Peptide elution was monitored by UV absorbance at 215 and 280 nm. Fractions were pooled across the UV elution profile to give 12 fractions for LC-MS/MS acquisition. Peptide fractions were dried in a vacuum concentrator before reconstituting in 20 ml aqueous 0.1% (v:v) trifluoroacetic acid (TFA).

TMT labeled peptides fractions were loaded onto an mClass nanoflow UPLC system (Waters) equipped with a nanoEaze M/Z Symmetry 100 Å C_18_, 5 µm trap column (180 µm x 20 mm, Waters) and a PepMap, 2 µm, 100 Å, C_18_ EasyNano nanocapillary column (75 mm x 500 mm, Thermo). The trap wash solvent was aqueous 0.05% (v:v) trifluoroacetic acid and the trapping flow rate was 15 µL/min. The trap was washed for 5 min before switching flow to the capillary column. Separation used gradient elution of two solvents: solvent A, aqueous 0.1% (v:v) formic acid; solvent B, acetonitrile containing 0.1% (v:v) formic acid. The flow rate for the capillary column was 330 nL/min and the column temperature was 40°C. The linear multi-step gradient profile was: 2.5-10% B over 10 mins, 10-35% B over 75 mins, 35-99% B over 15 mins and then proceeded to wash with 99% solvent B for 5 min. The column was returned to initial conditions and re-equilibrated for 15 min before subsequent injections. The nanoLC system was interfaced to an Orbitrap Fusion hybrid mass spectrometer (Thermo) with an EasyNano ionization source (Thermo). Positive ESI-MS, MS^2^ and MS^3^ spectra were acquired using Xcalibur software (version 4.0, Thermo). Instrument source settings were: ion spray voltage, 2,100 V; sweep gas, 0 Arb; ion transfer tube temperature; 275°C. MS^1^ spectra were acquired in the Orbitrap with: 120,000 resolution, scan range: *m/z* 380-1,500; AGC target, 2e^5^; max fill time, 50 ms. Data dependent acquisition was performed in top speed mode using a 4 s cycle, selecting the most intense precursors with charge states 2-6. Dynamic exclusion was performed for 50 s post precursor selection and a minimum threshold for fragmentation was set at 3e^4^. MS^2^ spectra were acquired in the linear ion trap with: scan rate, turbo; quadrupole isolation, 1.2 *m/z*; activation type, CID; activation energy: 35%; AGC target, 1e^4^; first mass, 120 *m/z*; max fill time, 35 ms. MS^3^ spectra were acquired in multi notch synchronous precursor mode (SPS^3^), selecting the 5 most intense MS^2^ fragment ions between 400-1,000 *m/z*. SPS^3^ spectra were measured in the Orbitrap mass analyser using: 50,000 resolution, quadrupole isolation, 1 *m/z*; activation type, HCD; collision energy, 65%; scan range: *m/z* 110-500; AGC target, 4e^5^; max fill time, 10 ms. Acquisitions were arranged by Xcalibur to inject ions for all available parallelizable time.

#### Protein identification and TMT label intensity quantification

Peak lists in .raw format were imported into PEAKS StudioX Pro (Version 10.6 Bioinformatics Solutions Inc.) for peak picking, database searching and relative quantification. MS2 peak lists were searched against 19,716 *Chlamydomonas* protein sequences appended with common proteomic contaminants. Search criteria specified: Enzyme, trypsin; Max missed cleavages, 1; Fixed modifications, TMT16plex (K and N-term peptide); Variable modifications, Oxidation (M); Peptide tolerance, 3 ppm; MS/MS tolerance, 0.5 Da; Instrument, ESI-TRAP. Peptide identifications were filtered to achieve a 1% peptide spectral match false discovery rate as assessed empirically against a reversed database search. Protein identifications were further filtered to require a minimum of two unique peptides per protein. TMT reporter ion intensities acting as markers of relative inter-sample peptide abundance were extracted from MS^3^ spectra for quantitative comparison. Protein level quantification significance used ANOVA for multi-way comparison and the PEAKSQ significance test for binary comparisons. In both cases the null hypothesis was that individual protein abundance was equal between groups. Normalization of label intensity was then carried out using the global ratio derived from total intensity of all labels. The Fold changes between comparison groupings were calculated based on their normalized TMT reporter ion intensities. Proteins which were not detected in all replicates for an individual bait were removed from calculation. Missing values were then replaced by the minimal value detected from each bait. Significance was determined via PEAKSQ test represented as - Log_10_P value.

### Recombineering cloning for localisation

Cloning of fluorescent protein tagged constructs was performed as previously described (Emrich-Mills et al., 2021). Briefly, homology arms to target genes at the 5’ of the native promoter and 3’ UTR were added to destination vectors via PCR. Homology arms of Cre13.g573250 were cloned into the pLM162-mScarlet-I backbone. Homology arms of Cre16.g663150, Cre06.g271850, Cre03.g172700, Cre17.g720450, Cre09.g394510, Cre03.g158050 and Cre02.g093650 were cloned into the pLM099-Venus backbone. Amplified backbones were transformed by electroporation into *E. coli* containing a Bacterial artificial chromosome and RecA vector which drives the recombination event. Resulting plasmids were selected on LB-agar plates containing Kanamycin and junctions confirmed by sequencing.

### Imaging of fluorescently tagged lines

For imaging of fluorescently tagged lines, photoautotrophically grown cells were immobilized on 1.5% low melting point agarose in TP-medium. Indirect immunofluorescence of RBCS2-APEX2 was performed according to (Uniacke et al., 2011) with the following modifications: Cells were fixed with 3.7% formaldehyde solution in PBS for 30 minutes at room temperature.

Anti-Flag antibody (F1804; Sigma-Aldrich) at 1:1000 dilution in PBS supplemented with 1% BSA was used as primary antibody. Anti-Mouse Alexa Fluor plus 555 (A32727; Invitrogen) was used as the secondary antibody at 1:1000 dilution. Labeled cells were then kept in the dark prior to imaging. Images were taken using a Zeiss LSM880 microscope with the Airyscan module or a Zeiss Elyra7 Lattice SIM. Excitation and emission filters of fluorophore and chlorophyll autofluorescence were set as follows: mVenus (Excitation: 514 nm, Emission: 520-550 nm); Chlorophyll (Excitation: 633, Emission: 610-650 nm); mCherry/mScarlet-I/Alexa Fluor plus 555 (Excitation: 561 nm, Emission: 580-600 nm).

### Amplex-red assay

Amplex™ UltraRed assay for RBCS2-APEX2 peroxidase activity was carried out according to the manufacturer’ s manual. Briefly, Amplex-red reagent (Fisher scientific; Invitrogen™ Amplex™ UltraRed Reagent #10737474) was dissolved in DMSO to a 10 mM stock. RBCS2-APEX2 and the untagged WT were grown photoautotrophically and split into triplicates. Cells were then chilled on ice for 5 minutes before resuspending in 200µL of reaction buffer (50 M Amplex Red, 2 mM H_2_O_2_ in PBS pH 7.4). The reaction was carried on ice for 15 minutes. Resorufin fluorescence measurement was performed using a Clariostar Plus Microplate reader using the following excitation and emission settings: Resorufin (Excitation: 535-555 nm; Emission: 580-620 nm); Chlorophyll autofluorescence (Excitation: 610-630 nm; Emission: 660-695 nm).

## Acknowledgements

C.S.L was supported by the Bill and Melinda Gates Foundation (Investment ID 53197). The project was supported by a United Kingdom Research and Innovation Future Leaders Fellowship to L.C.M.M (MR/T020679/1), Biotechnology and Biological Sciences Research Council Grants (BB/T017589/1, BB/S015337/1, BB/R001014/1, BB/X003035/1) and Engineering and Physical Research Council Grant (EP/W024063/1). P.G was supported by the Deutsche Forschungsgemeinschaft (DFG, German Research Foundation) – project number 456013262. The York Centre of Excellence in Mass Spectrometry was created thanks to a major capital investment through Science City York, supported by Yorkshire Forward with funds from the Northern Way Initiative, and subsequent support from EPSRC (EP/K039660/1; EP/M028127/1). The authors would like to thank the University of York Biosciences Technology Facility for confocal microscopy access and support and Masa Onishi for constructive discussions during the establishment of TurboID.

## Author Contributions

L.C.M.M. guided and supervised the project; C.S.L. designed and performed the biotin labeling experiments; A.D. performed LC-MS/MS analysis; C.S.L. and P.G. performed the fluorescent protein tagging and confocal imaging. C.S.L. and L.C.M.M. analyzed the data and wrote the manuscript with contributions from P.G., G.H.T. and A.D.; All authors discussed the results and commented on the manuscript.

## Supplementary Figures

**Supplemental Figure 1.**
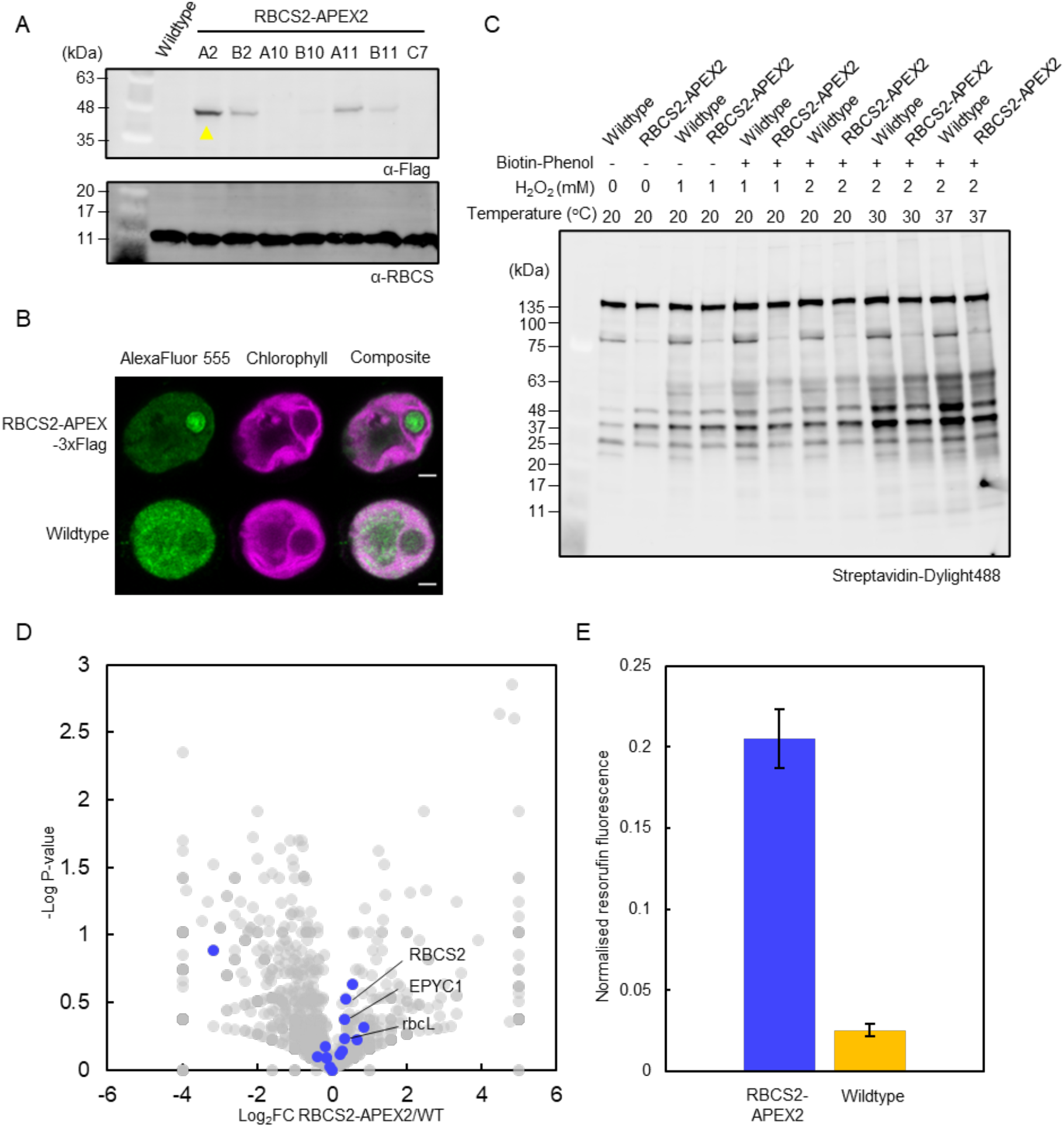
APEX2 does not efficiently label pyrenoid proteins in *Chlamydomonas* plastid. **A**, expression of RBCS2-APEX2 in *Chlamydomonas* CC-4533 was verified with immunoblotting the whole cell lysate with anti-HA. Anti-RBCS was used as loading control. Yellow arrow denotes the strain chosen for later labeling experiments. **B**, Localization of the RBCS2-APEX2 fusion protein was determined by immunofluorescence using anti-3xFlag. Green and Magenta signals denote the AlexaFluor555 and chlorophyll fluorescence respectively. Scale bar: 2 µm. **C**, Labeling efficiency of RBCS2-APEX2 was tested by incubating expressing cell lines with Biotin-phenol substrate at 2.5 mM for 2 hours. H_2_O_2_ activator was added at different concentrations (0 - 2 mM) and activation was carried out at a range of temperature (20 - 37°C). Biotin labeling was visualized by immunoblotting whole cell lysate against Streptavidin. **D**, Volcano plot representing the Log2 Fold Change of spectra count from RBCS2-APEX2 compared to untagged WT. Grey and dark blue dots represent detected proteins and known pyrenoid proteins respectively. Significance was determined via T-test. Proteins detected only in WT are set to -4 Log_2_ FC and proteins only detected in RBCS2-APEX2 are set to 5 Log_2_ FC. **E**, Amplex-red Assay was carried out to determine the peroxidase activity of RBCS2-APEX2 expressing lines. Untagged background and expressing cells were incubated with Amplex-red reagent and activated with H_2_O_2_ at 1 mM, the resorufin fluorescence emission from (Excitation: 535 - 555 nm; Emission: 580 - 620 nm) was normalized against chlorophyll autofluorescence (Excitation: 610-630 nm; Emission: 660-695 nm) (n=6).

**Supplemental Figure 2.**
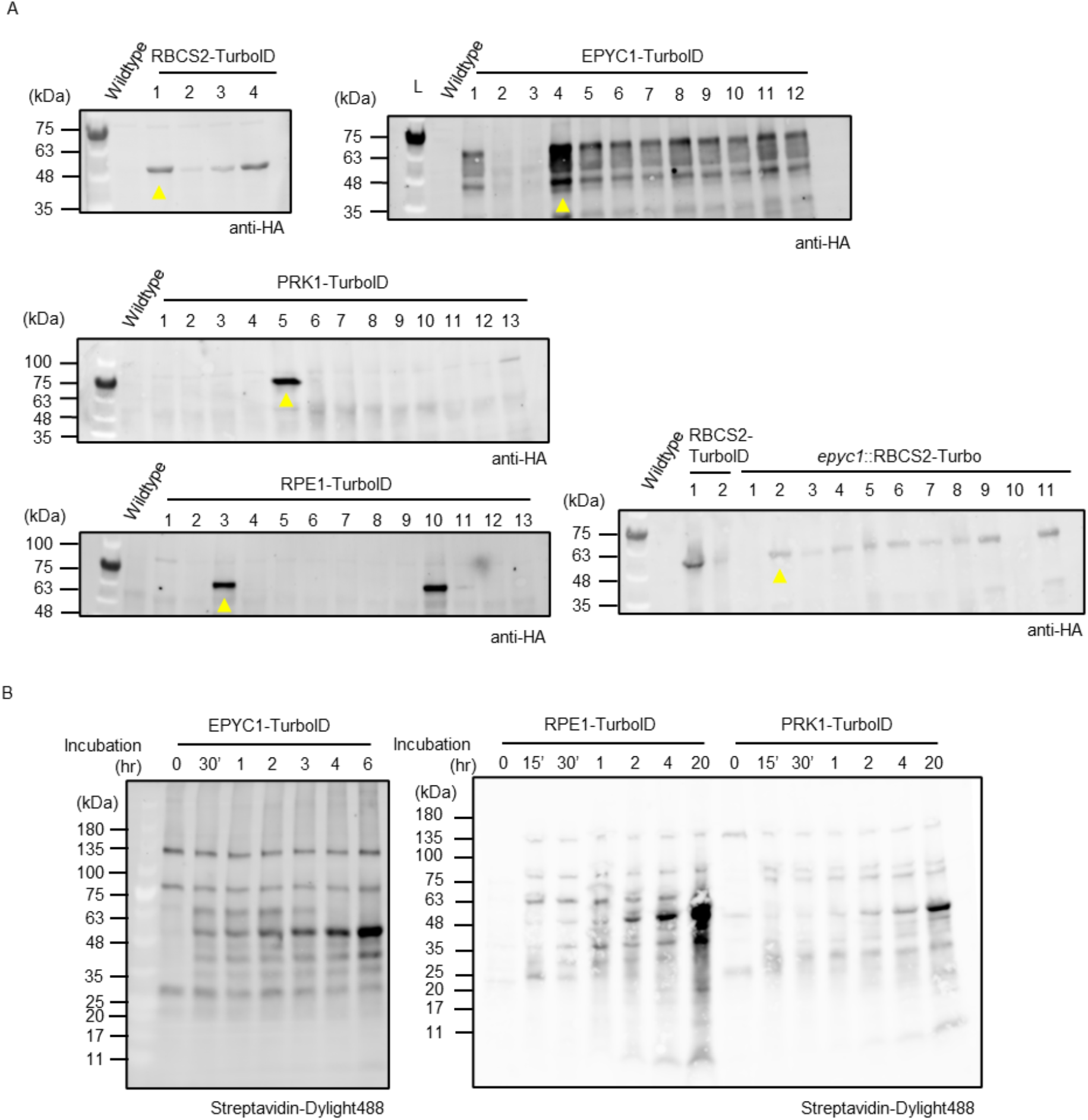
Screening strains for TurboID expression and activity. **A**, *Chlamydomonas* cells transformed with TurboID-tag plasmids were grown in TAP medium. Expression was assessed via Immunoblotting the whole cell lysate of picked strains with anti-HA antibody. Yellow arrows denote the strains used in labeling experiments in Figures 2, 3 and 5. **B**, Labeling activity of the various TurboID-tagged lines were assessed by incubating them in 2.5 mM biotin for a range of incubation time (0-20 hours).

**Supplemental Figure 3.**
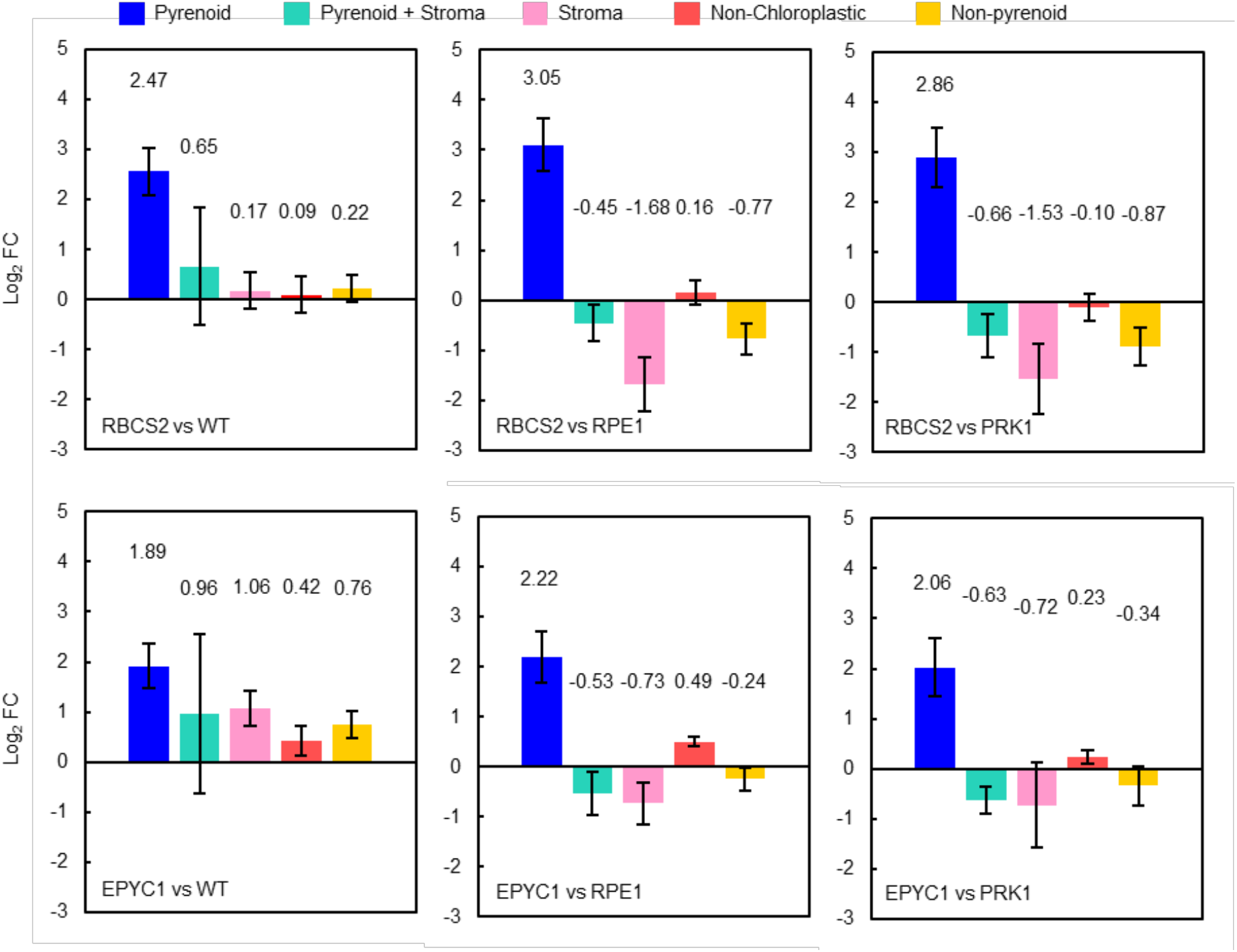
Enrichment of differentially localized proteins found in Figure 3. Averaged Log2FC for each comparison group was calculated according to their localization classification, a final category “non-pyrenoid” was created by combining all the non-pyrenoid proteins. Pyrenoids (n=16-19), Pyrenoid+Stroma (n=3-5), Stroma (n=9-13) and Non-chloroplastic (n=10), non-pyrenoid (n=23-27).

**Supplemental Figure 4.**
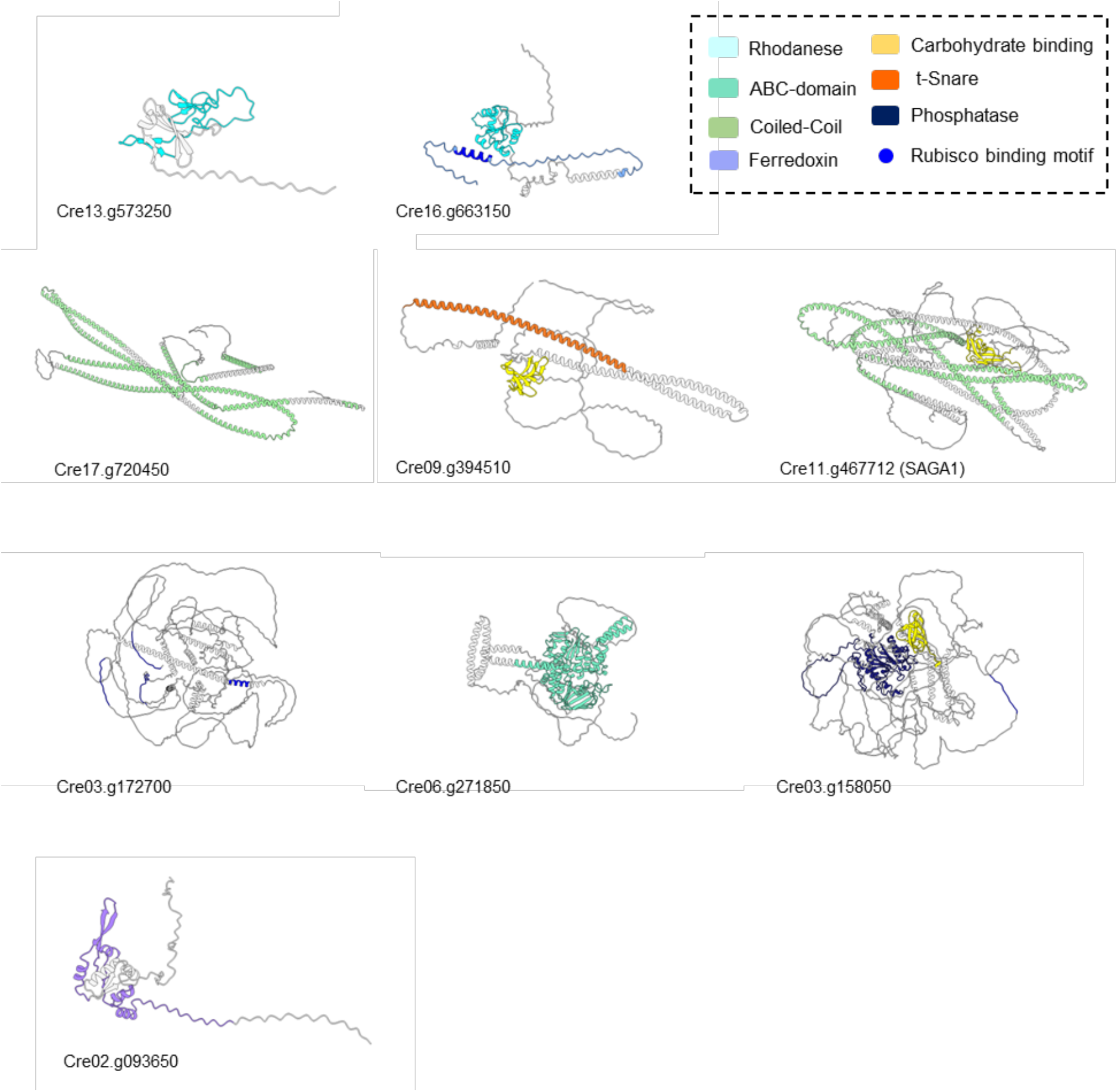
AlphaFold modeled structures for the uncharacterized proteins localized in this study. Structure models were obtained from Uniparc archive (UniProt Consortium, 2021), or if unavailable submitted manually for AlphaFold modeling (Jumper et al., 2021). For those manual submissions, modeling used the full preset db, with a max templating date of 2022-11-22. 5 models were completed and the number 1 ranked model based on pLDDT score is presented. Color scheme of predicted structures is listed which follows that of Figure 4B.

